# SINGLe: Accurate detection of single nucleotide polymorphisms using nanopore sequencing in gene libraries

**DOI:** 10.1101/2020.03.25.007146

**Authors:** Espada Rocío, Zarevski Nikola, Dramé-Maigné Adèle, Rondelez Yannick

**Affiliations:** Gulliver, ESPCI Paris, PSL University, CNRS, 75005 Paris, France

**Keywords:** nanopore sequencing, SNP detection, gene library, next generation sequencing

## Abstract

Nanopore sequencing is a powerful single molecule DNA sequencing technology which offers high throughput and long sequence reads. Nevertheless, its high native error rate limits the direct detection of point mutations in individual reads of amplicon libraries, as these mutations are difficult to distinguish from the sequencing noise.

In this work, we developed SINGLe (SNPs In Nanopore reads of Gene Libraries), a computational method to reduce the noise in nanopore reads of amplicons containing point variations. Our approach uses the fact that all reads are very similar to a wild type sequence, for which we experimentally characterize the position-specific systematic sequencing error pattern. We then use this information to reweight the confidence given to nucleotides that do not match the wild type in individual variant reads. We tested this method in a set of variants of KlenTaq, where the true mutation rate was well below the sequencing noise. SINGLe improves between 4 and 9 fold the signal to noise ratio, in comparison to the data returned by the basecaller guppy. Downstream, this approach improves variants clustering and consensus calling.

SINGLe is simple to implement and requires only a few thousands reads of the wild type sequence of interest, which can be easily obtained by multiplexing in a single minION run. It does not require any modification in the experimental protocol, it does not imply a large loss of sequencing throughput, and it can be incorporated downstream of standard basecalling.

## Background

Nanopore is a powerful technology for high throughput DNA sequencing, currently commercialized by Oxford Nanopore Technologies [1]. It provides sequence base calls reconstructed from conductivity records during the translocation of a single DNA molecule through a protein pore. This approach is implemented for example in a small device, minION, which offers portability and real time sequencing, using simple experimental protocols, and for a relatively low cost. A minION device can read DNA strands of various lengths, from PCR products up to megabase genomic fragments, and current versions return at least 5×10^9^ bases in one run. Therefore, it is an attractive device for sequencing libraries of amplicons that are too long for other next generation sequencing technologies. Unfortunately, nanopore’s relatively high error rate (≈ 5-10%) prevents the accurate detection of point genetic variation directly from individual reads [2,3]. Previous work aiming at high quality sequencing from nanopore data has concentrated on polishing tools such as Nanopolish [4], Racon [5] and Medaka [6]. These approaches start from a draft assembly and use the coverage depth to compute an averaged consensus at each position, via various computational approaches. Nanopolish, which uses the raw nanopore signal, reports an accuracy over 99% for a 29x sequencing coverage, while Racon reports 97% for 59x. Medaka reports 98% of accuracy in detection of single nucleotide polymorphisms (SNP) with a coverage of 100x. While these tools primarily apply to genome assembly, a number of experimental protocols were developed in order to leverage these pipelines in the specific case of amplicon library sequencing. These strategies aim to read -and associate- several replicates of the same molecule. This has been achieved by creating sequence concatenates using rolling circular amplification [7,8,9], which retrieved an accuracy of 99.5% for coverage of 150x, and via gene barcoding prior to amplification [10] with a reported accuracy over 99.9% for 25x coverage. Inconveniently, these methods add experimental efforts to the preparation of the sample prior to sequencing, are sensitive to bias occurring during the amplification steps [11], and reduce the number of different variants that can be studied, because a part of nanopore sequencing throughput is invested in reading each sequence many times.

In this paper, we propose a computational tool for amplicon sequencing, which applies in the case where the library is highly diverse but has low variability, i.e. it contains many different sequences differing from each other by only a few point mutations. This happens for example in directed evolution experiments, where the genetic libraries typically originate from a single ancestral sequence (the wild type) that has been submitted to limited randomization, for example using error-prone replication [12]. We propose a computational protocol, SINGLe (SNPs In Nanopore reads of Gene Libraries), which improves variant detection in individual reads from such libraries, for which a reference gene is known. In contrast to previous work, this approach uses standard 1D protocol minION sequencing and library preparation, and has a very limited impact on throughput.

We base our method on two observations, made during the sequencing of many identical copies of the reference sequence. First, the confidence or quality scores (Qscore) assigned by the base calling process to each nucleotide are usually low when a wrong nucleotide is assigned (supplementary figure S2), as expected. Second, the errors are not homogeneously distributed, and they are more frequent in some positions of the DNA (supplementary figure S1). These observations suggest that it should be possible to reduce the non-random part of the sequencing errors, using the information contained in the Qscore. The method we propose has two steps: the first one uses the reference reads to build a statistical model of the error pattern. Here we used a position and nucleotide-specific logistic regression. In the second step, this information is used to re-analyze base calls for the variant library and to update the confidence value of each nucleotide read in this dataset.

We tested SINGLe using the gene of KlenTaq DNA polymerase, a truncated variant of the well-known Taq polymerase, approximately 1.7 kb in length. We trained our model using approximately 6000 reads of the wild type and applied it on a toy library containing 7 known variants with 2 to 8 point mutations. Our computational correction reduced the sequencing noise, allowing a better identification of true point mutations. The signal to noise ratio is between 3 and 9 times better than the analysis performed directly over the guppy basecaller results. To illustrate the utility of this improvement, the corrected confidence values were used to classify sequences read by minION according to their sequence mutations: we reached an accuracy of 96% (versus 63% for guppy confidence values) leaving aside barcodes or other physical links. Furthermore, in the context of point mutant libraries, the corrected confidence values reduce the number of reads needed to obtain an exact consensus sequence by at least 5 times, with good performance obtained using only typically 10 reads. We show that this correction also outperforms the use of Racon and Nanopolish, state-of-the-art tools for consensus computation of nanopore sequencing reads of genome fragments. Finally, we schematize a possible pipeline for gene library analysis downstream SINGLe. Our results suggest that it is possible to accurately extract all variants within a library with a depth coverage of 20x, without using any physical links or barcodes between the identical strands.

## Method

Like other sequencing approaches, nanopore data analysis pipeline provides reads where each base is associated with a confidence value (Qscore). This number reflects the probability that the assigned nucleotide at that position is the correct one *p*_*right*_, via 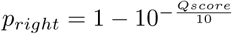.

We first characterized the errors and Qscore distributions on more than 6000 reads of the wild type KlenTaq gene (length 1665 nucleotides), for which we have a ground truth sequence (see supplementary table S1). In this data set, we can confidently attribute mismatches between the read nucleotide and the wild type as sequencing errors. At each position, and for each of the three non-wild type nucleotides and deletions, we quantified the proportion of errors made by guppy basecaller or, equivalently, the proportion of correct reads (which we denominate *n*_*correct*_), and grouped them according to the Qscore provided by the basecaller. *n*_*correct*_ is generally smaller for low Qscore (figure 1A and S4), reflecting that most errors appear where guppy base-calling confidence is low (figure 1C). However, the pattern strongly differs base-to-base and position-to-position (see examples in supplementary figure S4). We fitted this relation position and base-wise by a logistic regression, which provided a classifier able to convert the reported Qscore to the probability that this read is indeed correct (*p*_*right*_). This was made separately for the sequences read for sense and antisense strands.

**Figure 1:**
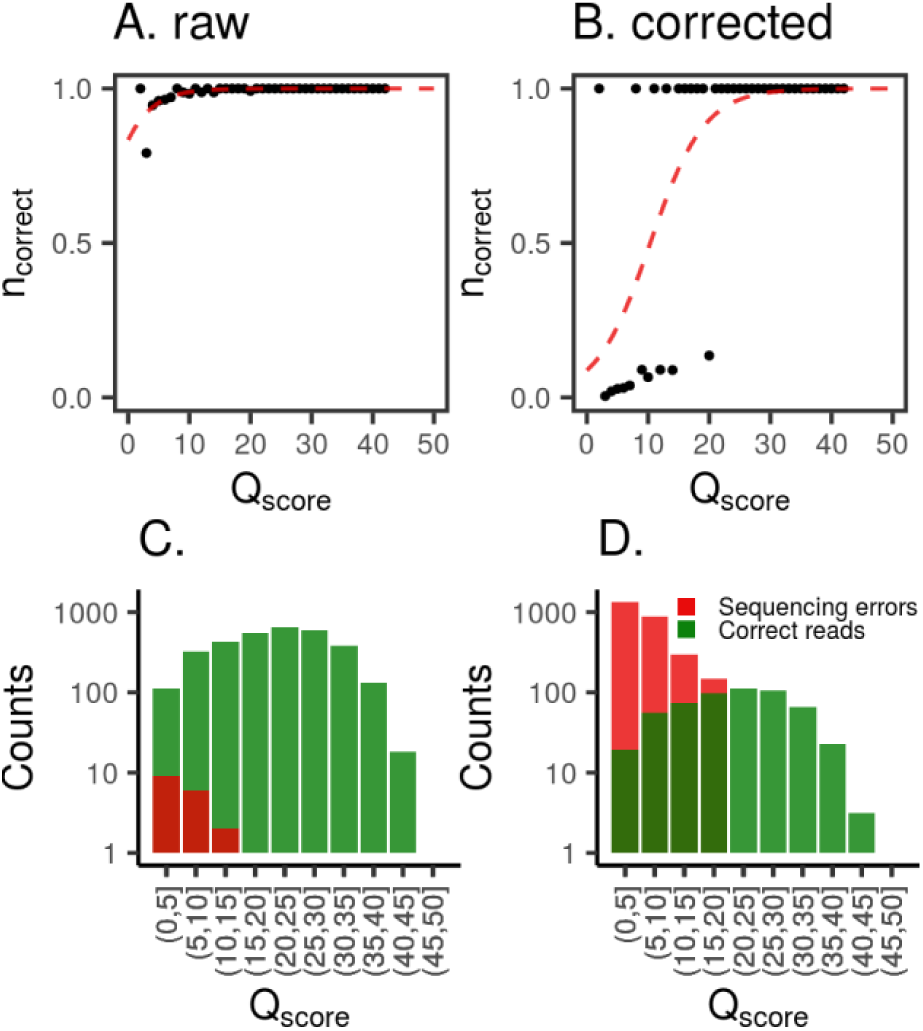
Example of logistic regression over reads of a known wild type sequence. A: black dots are the proportion of correct reads against the Qscore reported by guppy base-caller. Red line is the logistic regression performed over this data. C: Distribution of correct reads (green) and sequencing errors (red). B and D: Same plots as A and C, when each read is weighted according to the prior probability of actual mutation.

To include deletions in this analysis (which do not have a Qscore assigned by the basecaller), we fixed their confidence value as the average of the Qscore of their nearest neighbors in the nucleotide sequence. This decision was inspired by the observation that Qscores are correlated between consecutive nucleotides (supplementary figure S3). Insertions were ignored. All together, we obtained 13320=1665×4×2 regressions, one for each position of the gene (1665 bp), for each non-wild type nucleotide or deletion possible (4 possibilities in total for each position) and for the forward and reverse sense of sequencing.

With this information in hand, we then looked at mismatches in reads originating from mutated variant sequences. For each of them, we used the corresponding logistic regression to compute *p*_*right*_, the probability of actually being a mutation according to the associated Qscore. The higher the value of *p*_*right*_, the more likely it is a true mutation. However, the training set uses wild type sequences, which do not contain true mutations and hence our naive classifier is heavily biased against classifying mutations, as represented in figures 1A and 1C. It thus discards almost all mutations, even if they were correctly read by the nanopore sequencer. This is not representative of the actual proportion of errors/correct reads present in the mismatches of a set of mutants. To adapt the classifier, we need an a priori expectation of mutations (*p*_*prior*−*right*_), which must come from independent information. In the present case, the variant sequences originate from an error prone PCR (ePCR) process, for which we possess an estimate of the mutation frequency. We also obtain the a priori expectation of an observed mismatch to be a sequencing error (*p*_*prior*−*error*_) as the sequencing error rate at that position in the wild type set. We therefore compute *p*_*prior*−*right*_ and *p*_*prior*−*error*_, which we use to reweight the wild type set before fitting. This shifts the logistic regression towards the higher Qscore, allowing the classifier to accept a number of observed mismatches consistent with the prior expectation (figure 1B and D). We then proceed to re-score the mismatches observed in the mutant library, taking into account position, nucleotide and sense of the strand.

## Results

To test our method, we built a toy library of KlenTaq mutants and sequenced them using the minION device (see experimental methods). The library consists of 7 known variants uniquely barcoded before sequencing, thus we know exactly what mutations to expect in each read. The complete list of mutations is available in supplementary table S2. We used the logistic regressions performed on the wild type data to convert Qscore into a probability of being a true mutation the *p*_*right*_ of each nucleotide, as described in section Method.

### Signal to noise ratio

The high error rate in Nanopore sequencing makes it difficult to distinguish actual mutations from sequencing errors in single reads. A straightforward procedure to filter errors is to only trust read positions which have a high probability of being correct. In this section, we compare how this threshold process performs when using, either, the raw *p*_*right*_ returned by guppy basecaller on nanopore reads, the *p*_*right*_ after fitting with a logistic regression (or correction naive), or *p*_*right*_ after fitting with a logistic regression using prior probabilities (SINGLe).

We used the toy library to quantify the signal to noise ratio. In this example each sequence is barcoded, so we know if each mismatch is a sequencing error or an actual mutation. We defined signal as the number of mismatches known to be mutations with a *p*_*right*_ higher than the threshold (true positives), and noise as the number of those mismatches known to be non-mutated positions (false positives). The counts are weighted by *p*_*right*_ for each nucleotide. Results are shown in figure 2. For all thresholds, SINGLe has a higher signal to noise ratio (between 4 and 9 times higher, depending on the cut off), thus facilitating the identification of actual mutations. This remains true when no cut off is applied (cut off = 0). Results are similar if we count the number of mismatches over the threshold without weighting them by *p*_*right*_ (supplementary figure S5). We noticed that the signal to noise ratio improvement is mainly driven by the reduction of the false positive rate, i.e. reduction of sequencing noise. Nevertheless, the true positive rate also decreases fast (supplementary figure S6). This should be taken into account if the goal is to detect mutations which are poorly represented in the sequenced set.

**Figure 2:**
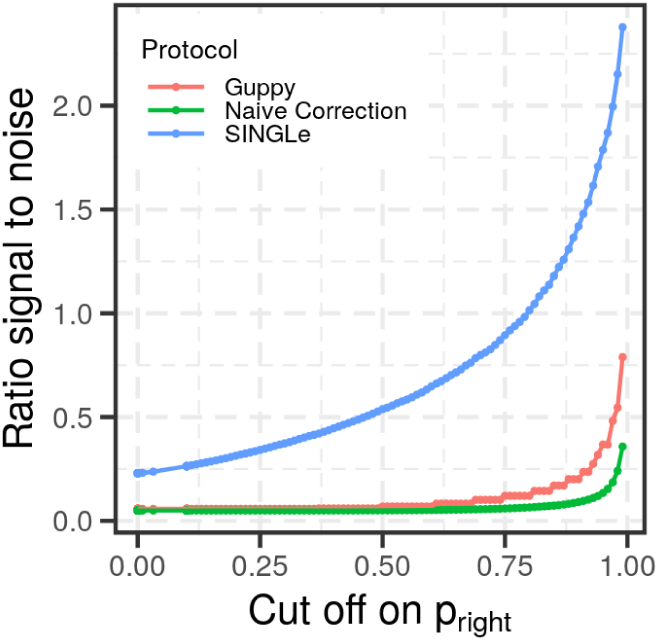
Signal to noise ratio as a function of the cut-off on the quality values *p*_*right*_, for the three methods analyzed: using *p*_*right*_ provided by guppy basecaller (red), using *p*_*right*_ after naive correction (green) and *p*_*right*_ returned by SINGLe correction with priors (blue). SINGLe improves the signal to noise ratio for any *p*_*right*_ cut off.

### Sequences clustering

To show that the reduction of noise is relevant to characterize an ensemble of reads into sequence clusters, we used the corrected probabilities *p*_*right*_ to group reads by weighted sequence similarity. We defined the dissimilarity between two reads s1 and s2 as a modified Hamming distance:

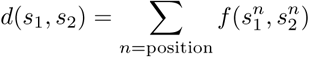

Where

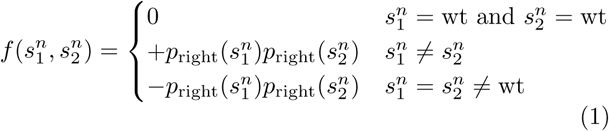

where *p*_*right*_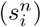 is *p*_*right*_ assigned to sequence i at position n. In other words, we sum zero if both sequences are equal to wild type (wt), sum the product of corrected probabilities of both sequences if they are different (one mutation present in only one of them), and subtracted the product of corrected probabilities of both sequences if they are the same and differ from wild type (both reads display the same mutation). Then we used R’s implementation of Ward’s minimum variance method to cluster the reads. We evaluated this procedure on the toy library of 7 variants. The resulting dendrogram is displayed in figure 3A, where each leaf is a different read, and the color of the point below represents which KlenTaq variant it is (according to its barcode). When cutting the dendrogram at a fixed height to obtain seven clusters, we can find a prevailing variant on each of them, which allows us to use this as a classification method. Out of 3538 reads analyzed, 3381 were correctly clustered (96%). This was not true when using guppy base-caller’s *p*_*right*_ values, in which only 63% were correctly classified (figure 3B).

**Figure 3:**
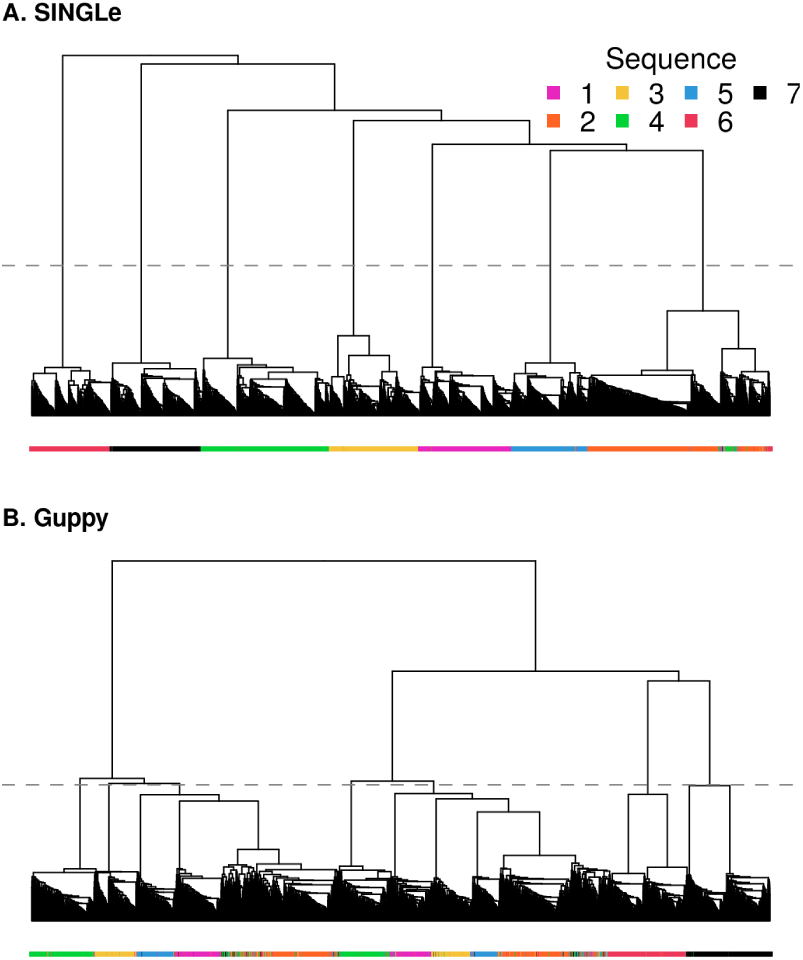
Hierarchical clustering of reads of a toy library constituted by 7 mutants of the KlenTaq gene, each of them experimentally identified by a barcode. The colors below each leaf indicate to which variant the read corresponds to. In A, distances between sequences were computed using Qscores by SINGLe, while in B we used the Qscores returned by guppy basecaller.

### Consensus calculation

One strategy for lowering noise in nanopore sequencing consists in computing a variant consensus sequence (VCS), i.e. reading several times a variant and using these reads to calculate the most likely sequence [4,11]. Usually, average on more reads leads to a more accurate VCS. We hypothesized that the corrected weights returned by SINGLe should help to converge faster on the actual sequence, as they increase the confidence on mutations over the sequencing errors. We tested this hypothesis over the set of 7 known KlenTaq variants sequenced using minION and basecalled using guppy.

We computed the VCS over a set of aligned reads of the same KlenTaq variant (using the fact that these reads were barcoded). For each position, we calculated the weighted frequency of each nucleotide and deletions. The weights used are the Qscore returned by guppy base-caller, or the ones returned by SINGLe. The nucleotide (or deletion) with the highest weighted count is kept as the consensus in that position.

We computed the consensus on a set of 2 to 50 sequences drawn randomly, and repeated 100 times for each set size. In figures 4 and S8 (red curves) we show how many times the VCS matches exactly the actual sequence. In all cases where there are no deletions (variants 1-4 and 7), or where there is a deletion in a non-homopolymer position (variant 6), the convergence on the actual sequence is faster when using SINGLe weights: perfect consensuses are obtained for more than 95% of attempts starting from typically 10-20 sequences. This performance is not reached in sets of up to 50 sequences by the Qscore of guppy basecaller.

**Figure 4:**
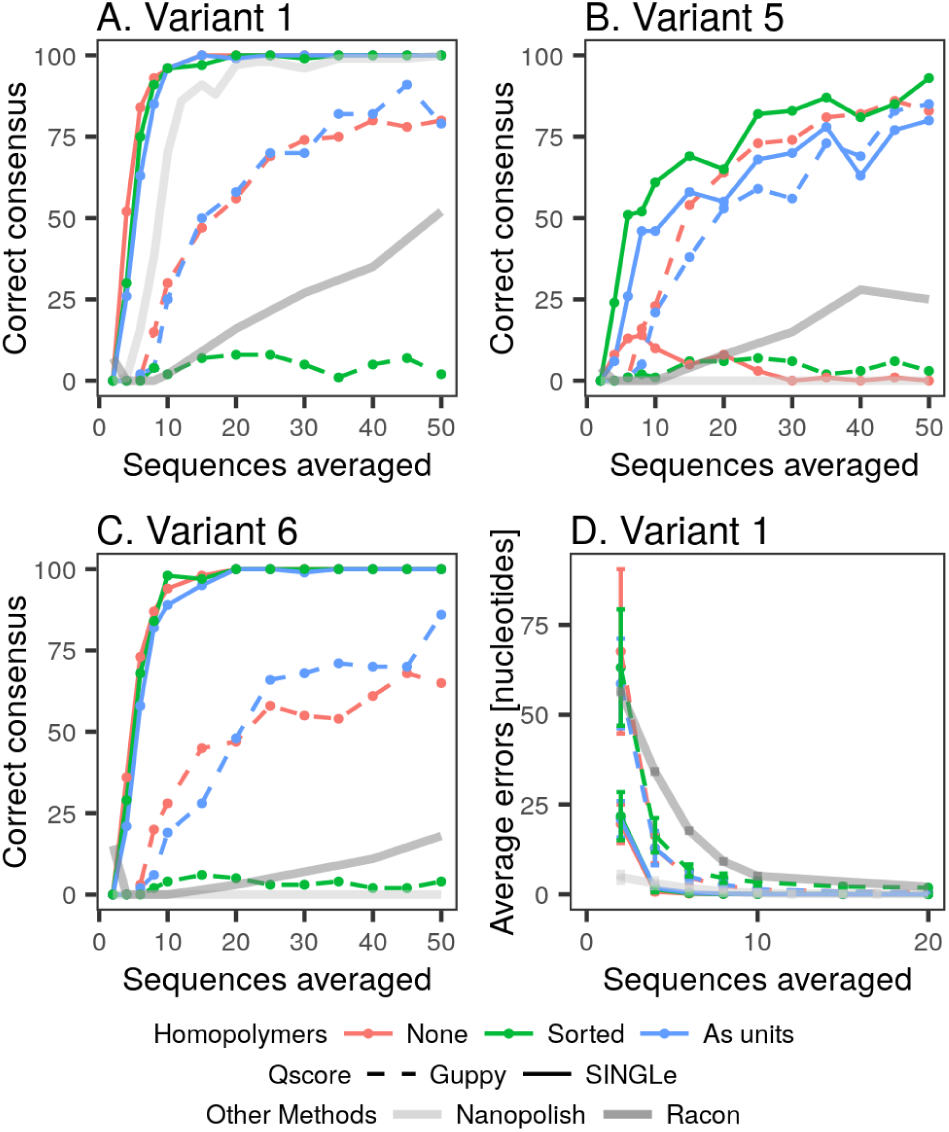
A-C Performance of consensus calling according to the number of reads used to compute it. Different strategies are used to compute the consensus: In red, no pre-processing of sequences. In green, sorting homopolymers to move deletions towards the end. In blue, treating homopolymers as a unit. Panel A shows the results for variant 1 (no deletions), panel B for variant 5 (which has a true deletion within a homopolymer), and panel C for variant 6 (which has one deletion in a non-homopolymeric position). Panel D shows the average error per consensus sequence according to the number of sequences used. In all cases we compare the use of guppy basecaller weights (dashed line) and weights corrected by SINGLe (solid line). In dark gray, results using Racon software, and in light gray using Nanopolish.

Results are different for variant 5, which has a deletion within a homopolymer (‘GG’ at positions 489 and 490), making it a more challenging target. The basic VCS matches performed worse with SINGLe scores and including more sequences even reduces the correct percentage (figure 4B, red curves). We noticed that this is produced by the misrepresentation of the deletion. The VCS is computed position by position, and the aligner used (LAST) does not always represent the deletion in the same position (sometimes is ‘G-’ and sometimes is ‘-G’). Thus the presence of a deletion is averaged out. We tested two strategies to address this issue: on the one hand we tried to sort all homopolymers so that the deletions are always at the end before computing the VCS. Results are shown in green curves in figure 4. In this case, the results for variant 5 improved, reaching a performance slightly better than using Qscore provided by guppy basecaller, while similar performance for all other variants compared to the basic consensus calculation. Notably, this strategy is counter-productive for the VCS calculation using guppy raw scores in all variants (green dashed lines). On the other hand, we tried evaluating the homopolymers as a unit instead of position by position thus being equivalent ‘G-’ to ‘-G’ (blue curves in figure 4). Once more, the results for variant 5 improved, reaching a performance as good as using Qscore from guppy base-caller, without affecting other variants. For a deeper discussion on this point, refer to the supplementary material.

We also evaluated how different the VCS are from true sequences. In figures 4D and S9, we plotted the mean number of wrong nucleotides in the VCS. When using the SINGLe weights, only a few replicas are necessary to recover the actual sequence with very few errors: 5 reads return a mean error below one nucleotide per VCS (except variant 5). This number is always over 10 sequences when using guppy raw Qscore as weights. Also, there are no significant changes among the different treatments of homopolymers in the consensus calculation.

As a reference, we computed consensus sequences using Racon [5] and Nanopolish [4], state-of-the-art tools for consensus calling in Nanopore reads (gray curves in figures 4, S8 and S9). In all cases, we observed that the convergence for Racon and Nanopore require more reads than the methods presented in this manuscript. However, we note that they do not use a reference dataset of wild type sequences, which makes direct comparisons difficult.

### Clustering and consensus calling for a non-barcoded library

Finally, we simulated the situation in which a gene library is sequenced at some depth (but without barcodes) and the goal is to recover the exact sequence of each variant and their relative abundance in the sample. We mixed the reads of the 7 variants and removed the barcode identities, randomly picked 140 of them, to return an average 20 reads per variant, and applied the following sequence: we implemented SINGLe, computed the modified hamming distance (equation 1) between each pair of reads, clustered them hierarchically by Ward’s minimum variance method, we obtained the dendrogram in figure 5A, we cluster them in 7 groups as suggested by the within cluster sum of squares method (supplementary figure S11), and we computed the VCS for each cluster using the sorting procedure for homopolymers. The sequences constructed through this procedure match exactly the actual variants.

**Figure 5:**
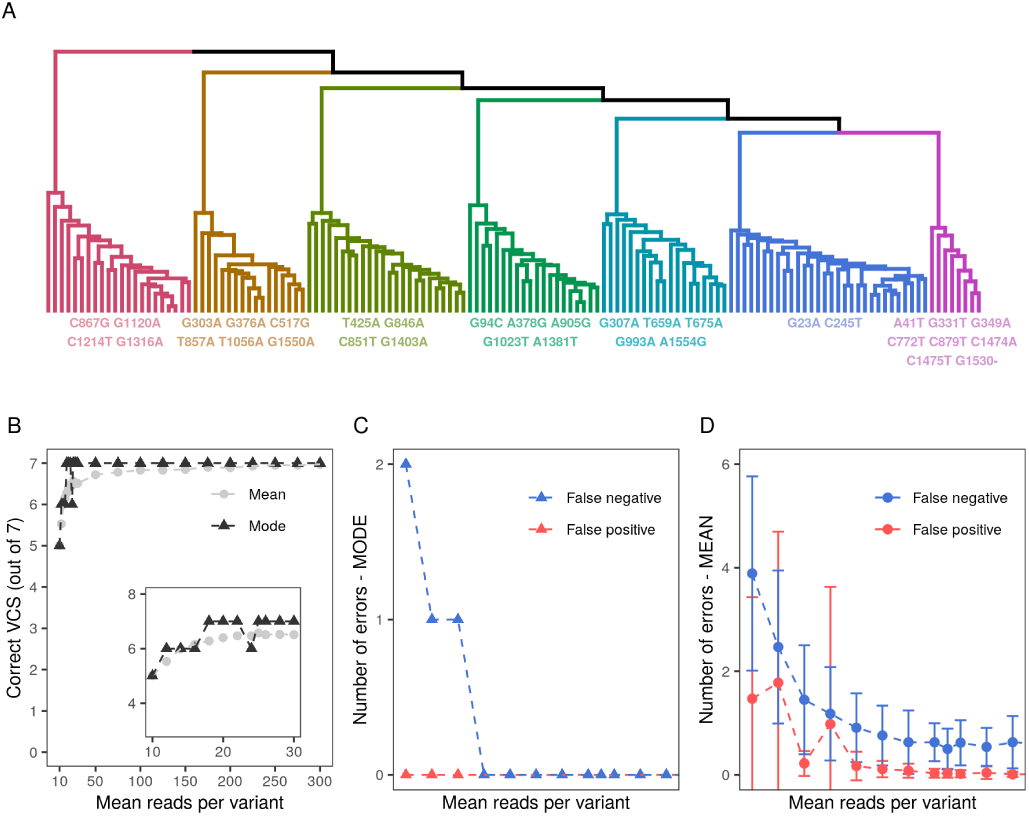
Cluster and consensus approach using SINGLe. The analysis was performed over a subset of randomly chosen reads. We corrected the Qscores using SINGLe, clustered the reads and computed the consensus sequence for each group, following the methods described in this manuscript. A. Dendrogram of reads, colored by cluster, and below the consensus sequence computed over 140 reads (20 per variant). B-D Robustness of the analysis according to the sequencing depth. B. Mean and mode of how many sequences out of the true seven variants are predicted with no mistakes. C-D. Out of the 35 mutations present in the 7 variant set, mode and mean of how many of them are not detected (false negatives), and how many predicted mutations are not in the original dataset (false positives).

To evaluate the robustness of this analysis, we repeated it for subsets of 70 to 2100 nanopore reads, i.e. an average of 10 to 300 reads per variant, with a bootstrapping of 100 replicas. We clusterized these sets in 7 groups and counted the number of variants that were exactly retrieved by group-wise VCS. In figure 5B we show the mode and average for each set size. Both the mean (equal to 6.5) and the mode (equal to 7) suggest that using a coverage of less than 20x is enough to recover the actual sequences from nanopore reads. This is confirmed by looking at false positives (non existing mutations appearing in the VCS) and false negatives (missed mutations) (figure 5C and 5D). Both mean and mode of these values indicate that false positives are rare for any set size and false negatives also quickly converge (with the tail being mostly due to the problematic homopolymer deletion in variant 5).

## Conclusions

The relatively high error rate in single molecule nanopore sequencing limits some applications such as the analysis of libraries containing many different, but genetically similar, sequences.. The approach we propose here, SINGLe, leverages the fact that the sequencing errors in this case are partly systematic, as previously noted [13]. We accumulate many reads from the reference gene to build a sequence-specific error model that can locally correct for the sequencing biases. Applying this procedure on a toy library of the KlenTaq gene with an average mutation rate of 3.3 bases/kb, we showed that correcting the confidence values provides a large increase in the signal to noise ratio. Consequently, the clustering of reads is improved to 96% accuracy (vs 63% using raw data), and consensus calling returns 95% of perfect results from typically 10-20 reads. An important ingredient of SINGLe is a correct prior for the number of mutations in the train and test set. Here, our reference sequence was assumed to be perfect, and we could evaluate precisely the average mutation rate in the test set, because it originated from a controlled experimental mutagenesis protocol. In other situations it would be possible to use short read sequencing, for example Illumina, to evaluate this number. If the full sequence is submitted to short read high quality sequencing, it would even be possible to obtain more precise priors, for example specific to each position and nucleotide. Our approach would then be used to phase these statistical mutations to single long reads.

An underlying assumption of our method is that the distribution of Qscore observed at a particular position for the wild type base reflects appropriately the distribution of Qscore that would be observed for a variant base at that position. This approximation is necessary since the error model is built from a single sequence and hence has a single ‘‘true” base per position. Fortunately, the difference in Qscore distributions for ‘‘true” versus ‘‘error” seems large enough for our method to perform well within that approximation.

SINGLe needs to characterize the sequencing errors done on an appropriate reference sequence. As such it is limited to analyze variants which are close neighbors of the reference, and where mutations can be considered to be independent. We did not try to adapt the method to detect alterations beyond point replacements or deletions, which may require a more complex analysis pipeline. Encouragingly, variant 6 contained two contiguous mutations, and was properly analyzed by our consensus approach. We also note that, in the case of highly diverse sequences, and if the library can be sequenced at sufficient depth, it becomes easier to cluster sequences and compute direct consensuses. Finally, to our knowledge SINGLe is the first tool fully focused in analyzing gene libraries sequenced by standard nanopore technology. Our approach provides large improvement of signal to noise at very little experimental effort or throughput reduction. This is because only the reference DNA needs to be sequenced many times, compared to other methods where each library member requires multiple reads. In SINGLe, the reference DNA dataset can be obtained simultaneously with the libraries, using standard barcoding protocols. There are no other modifications to the experimental protocol, and the computational error correction process can be simply added to any analysis pipeline after base calling. Therefore, in contrast to methods that increase accuracy at the cost of throughput, SINGLe paves the way to a more accurate characterization of large libraries of long genetic elements.

## Experimental methods

### Samples preparation

KlenTaq wild type gene was amplified using a high fidelity PCR (Q5 polymerase from NEB) from a stored plasmid. See DNA sequence in supplementary table S1.

Mutants were obtained via error prone PCR (ePCR) using Agilent’s kit GeneMorph II. We started from 1.1 nM of dam-methylated DNA. We used primers GGGATTATTCTTTGGCGCTCAGCCAAT and AC-CATGCGTCTGCTGCATGAAT. Thermocycling was performed as follows: 95C for 2 min, followed by 25 cycles of [95C for 30sec + 65C for 30sec + 72C for 2min] and a final extension at 72C for 10 min. We digested the product with DpnI (NEB) and purified it using columns (Macherey-Nagel). We put the mutagenized genes in a pIVEX vector via Gibson assembly (NEB Hi-Fi DNA assembly) using 125ng of gene DNA, 100 ng of vector in a 2:1 insert:vector molar ratio and incubated for 15min at 50C. We purified and concentrated DNA with a Zymo Research kit. We transformed the product into chemocompetent KRX bacteria. We spread them on a Petri dish with LB and Ampicillin. We incubated overnight and picked some clones randomly. We verified the presence of the plasmid via colony PCR (using DreamTaq polymerase from Thermofisher). 10 positive clones were grown overnight in liquid LB with antibiotic and mini-prepped to obtain the plasmid DNA. A fraction was used for high quality sequencing (Sanger sequencing), and another fraction used for minION sequencing.

### minION sequencing

We used the prepared DNA of each clone and the wild type gene for amplification by PCR with Q5 polymerase (NEB) using primers which included the minION barcodes adapters: ACTTGCCTGTCGCTC-TATCTTCAGTGTGCTGGAATTCGCCCTTTTA and TTTCTGTTGGTGCTGATATTGCAGACCA-CAACGGTTTCCCTCTAGAAATA. Thermocycling was performed as follows: 98C for 30sec, 23 cycles of [98C for 10sec + 59C for 30sec + 72C for 1min], final extension at 72C for 2min.

We digested the product with Dpn1 (NEB), gel purified it using Macherey-Nagel kit. We proceeded following standard Oxford Nanopore protocols for minION. We used one barcode for wild type and one for each of the mutants 1 to 7.We used kits EXP-PCB001 for barcoding, SQK-LSK108 for ligation, EXP-LLB001 for flow cell loading, and minION flow cell version was R9.4/FLO-MIN106, thus the sequencing was 1D.

### minION reads pre-processing

minION raw data was basecalled and demultiplexed using ONT Guppy version 3.3.3. We obtained 140197 reads from which 45635 (33%) did not match any barcode. Each sequence was pairwise aligned to wild type KlenTaq sequence using LAST [14]. Those sequences with low alignment score were discarded, keeping 32507 at the end (23% from original reads).

### Racon consensus

We selected a subset of sequences on the fastq file after base calling using an R script, aligned them to the wild type sequence using minimap2 2.17-r941 [15] (default parameters), and computed the VCS with racon v1.4.11 [5] (default parameters).

### Nanopolish consensus

We used Nanopolish 0.13.2. We selected a subset of sequences on the fasta file after base calling using an R script, and analysed them using the pipeline suggested by Nanopolish manual: ‘minimap2 -ax map-ont’ to align to reference, ‘samtools sort’ and ‘samtools index’ to sort and find indexes of reads and ‘nanopolish variants –consensus’ for consensus calling.

## Availability

Project name: SINGLe

Project home page: https://github.com/rocioespci/single.

Operating system: Linux

Programming language: R

Other requirements: awk

Licence: MIT

## List of abbreviations

ePCR: error-prone PCR
PCR: polymerase chain reaction
SINGLe: SNIPs In Nanopore reads of Gene Libraries
SNP: single nucleotide polymorphisms
Qscore: Quality score
VCS: Variant consensus sequence

## Funding

This project has received funding from the European Union’s Horizon 2020 research and innovation program under the Marie Sklodowska-Curie grant agreement No 845976 and from the European Research Council (ERC, Consolidator Grant No. 647275 ProFF).

## Competing interests

The authors declare that they have no competing interests.

## Notes

https://github.com/rocioespci/single

### Competing Interest Statement

The authors have declared no competing interest.

### Summary of Updates

- Renamed R package. - Updated the R package to make it simpler to use. - Improved pipeline for downstream analysis. - Improved benchmark.

