## Supplementary for "SINGLe: Accurate detection of single nucleotide polymorphisms using nanopore sequencing in gene libraries"

#### Sequence information

ATGCGTCTGCTGCATGAATTTGGTCTGCTGGAAAGCCCGAAAGCACTGGAAGAAGCCCCTTGGCCTCCGCCTGAAGGTGCATTTGTT
GGTTTTGTTCTGAGCCGTAAAGAACCGATGTGGGCTGATCTGCTGGCACTGGCAGCAGCACGTGGTGGTCGTGTTCATCGTGCACCG
GAACCGTATAAAGCACTGCGTGATCTGAAAGAAGCACGCGGACTGCTGGCAAAAGATCTGAGCGTTCTGGCCCTGCGTGAAGGTCTG
GGTCTGCCTCCGGGTGATGATCCGATGCTGCTGGCATATCTGCTGGATCCGAGCAATACCACACCGGAAGGTGTTGCACGTCGTTAT
GGTGGTGAATGGACCGAAGAAGCAGGCGAACGCGCAGCACTGAGCGAACGTCTGTTTGCAAATCTGTGGGGTCGTCTGGAAGGTGA
AGAACGTCTGCTGTGGCTGTATCGTGAAGTTGAACGTCCGCTGAGCGCAGTTCTGGCACACATGGAAGCAACCGGTGTTCGTCTGGA
TGTTGCCTATCTGCGTGCACTGAGCCTGGAAGTTGCAGAAGAAATTGCACGCCTGGAAGCAGAAGTTTTTCGTCTGGCAGGTCATCC
GTTTAATCTGAATAGCCGTGATCAGCTGGAACGTGTTCTGTTTGATGAACTGGGCCTGCCTGCAATTGGTAAAACCGAAAAAACCGG
TAAACGTAGCACCAGCGCAGCCGTTCTGGAAGCCCTGCGCGAAGCACATCCGATTGTTGAAAAAATTCTGCAGTATCGCGAACTGAC
CAAACTGAAAAGCACCTATATCGATCCGCTGCCGGATCTGATTCATCCGCGTACCGGTCGTCTGCATACCCGTTTTAATCAGACCGC
AACCGCCACCGGTCGCCTGAGCAGCAGCGATCCGAATCTGCAGAATATTCCGGTTCGTACACCGCTGGGTCAGCGTATTCGTCGTGC
ATTTATTGCCGAAGAAGGTTGGCTGCTGGTTGCACTGGATTATAGCCAGATTGAACTGCGTGTTCTGGCGCATCTGAGCGGTGATAA
AAATCTGATTCGTGTTTTTCAAGAGGGTCGCGATATTCATACCGAAACCGCAAGCTGGATGTTTGGTGTTCCGCGTGAAGCAGTTGA
TCCGCTGATGCGTCGTGCAGCAAAAACCATTAACTTTGGTGTGCTGTATGGTATGAGCGCACATCGTCTGAGCCAAGAACTGGCAAT
TCCGTATGAAGAAGCACAGGCCTTTATTGAACGTTATTTTCAGAGCTTTCCGAAAGTTCGTGCATGGCTGAAAAAAACCCTGGAAGA
GGGACGTCGTCGTGGTTATGTTGAAACCCTGTTTGGTCGTCGTCGCTATGTTCCGGATCTGGAAGCACGTGTTAAAAGCGTTCGTGA
AGCCGCAGAACGTATGGCCTTTAATATGCCGGTTCAGGGCACCGCAGCAGATCTGATGAAACTGGCCATGGTTAAACTGTTTCCACG
GCTGGAAGAAATGGGTGCACGTATGCTGCTGCAGGTTCATGATGAGCTGGTGCTGGAAGCGCCTAAAGAACGTGCAGAAGCCGTTG
CCCGTCTGGCCAAAGAAGTTATGGAAGGCGTTTATCCGCTGGCAGTTCCGCTGGAAGTGGAAGTTGGTATTGGTGAAGATTGGCTGA
GCGCCAAAGAATAA

**Table S1:** DNA sequence of the gene of Klen Taq wild type

| from | position | to | Variant ID |
|---|---|---|---|
| C | 867 | G | 1 |
| G | 1120 | A | 1 |
| C | 1214 | T | 1 |
| C | 1316 | A | 1 |
| G | 23 | A | 2 |
| C | 245 | T | 2 |
| G | 94 | C | 3 |
| A | 378 | G | 3 |
| A | 905 | G | 3 |
| G | 1023 | T | 3 |
| A | 1381 | T | 3 |
| T | 425 | A | 4 |
| G | 846 | A | 4 |
| C | 851 | T | 4 |
| G | 1403 | A | 4 |
| G | 307 | A | 5 |
| T | 659 | A | 5 |
| T | 675 | A | 5 |
| G | 993 | A | 5 |
| A | 1554 | G | 5 |
| G | 490 | - | 5 |
| A | 41 | T | 6 |
| G | 331 | T | 6 |
| G | 349 | A | 6 |
| C | 772 | T | 6 |
| C | 879 | T | 6 |
| C | 1474 | A | 6 |
| C | 1475 | T | 6 |
| G | 1530 | - | 6 |
| G | 303 | A | 7 |
| G | 376 | A | 7 |
| C | 517 | G | 7 |
| T | 857 | A | 7 |
| T | 1056 | A | 7 |
| G | 1550 | A | 7 |

**Table S2:** Point mutations identified in the 7 variants of Klen Taq in the toy library.

#### General observations

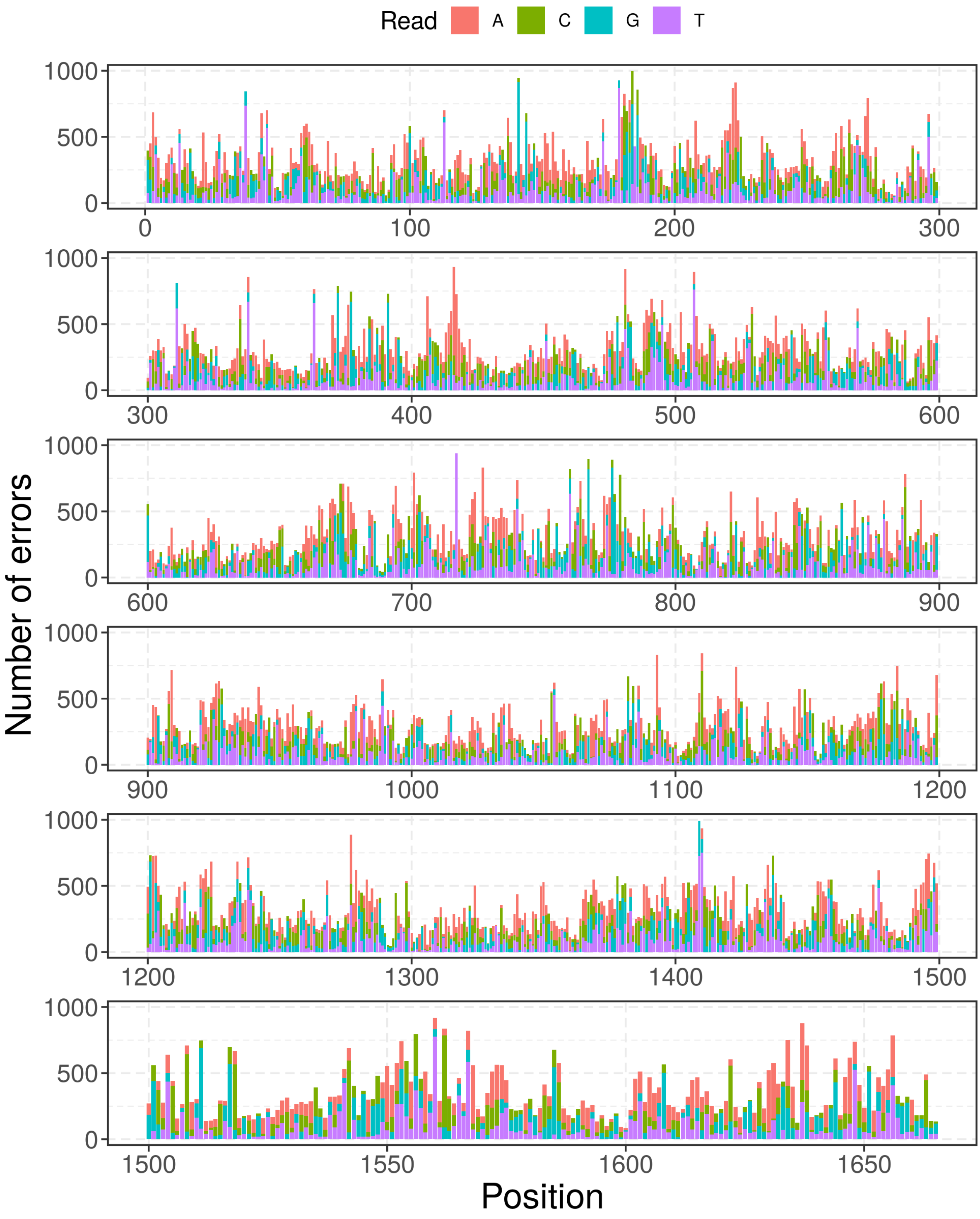


**Figure S1:** Errors made by minION when sequencing the wild type Klen Taq gene, classified by position (x axis) and nucleotide reported by the basecaller (color).

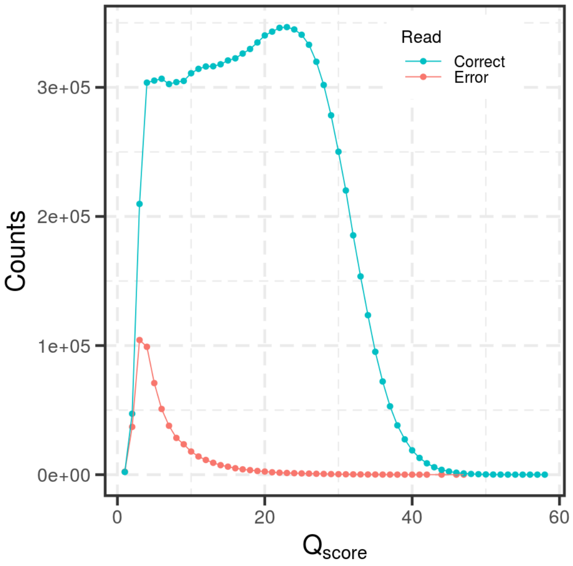


**Figure S2:** Distribution of nucleotide reads on the wild type sample according to the $Q_{score}$ assigned by minION and classified into correct reads (blue) or sequencing errors (red).

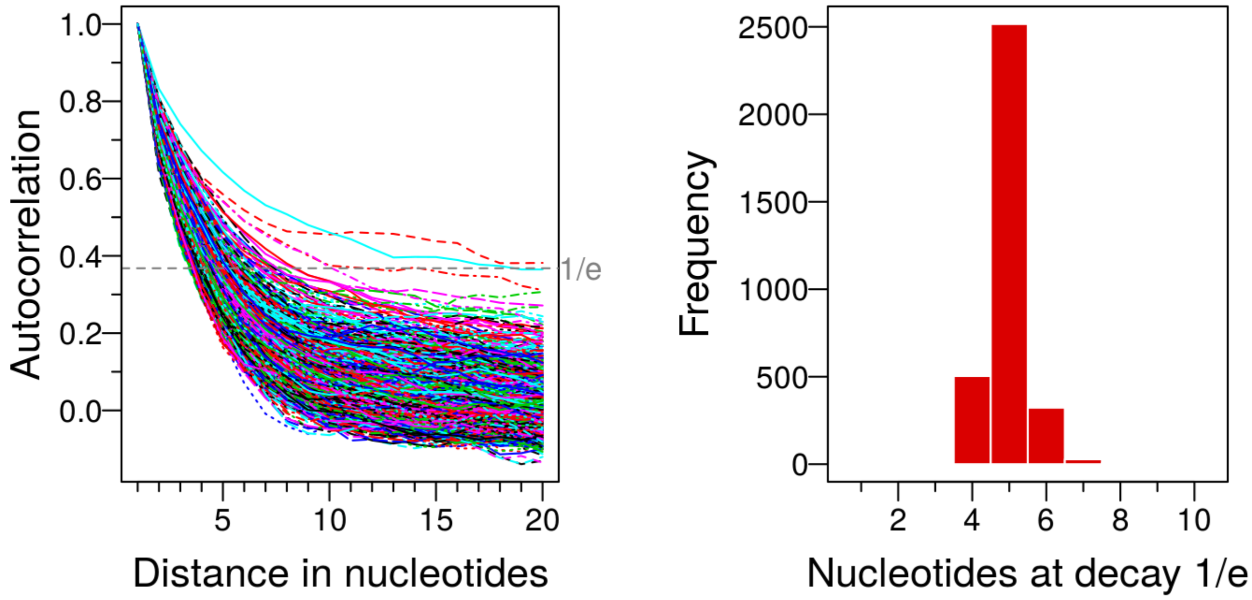


**Figure S3:** Left, autocorrelation function of the $Q_{score}$ on a DNA strand read by minION. Each line correspond to an independent read. All of them belong to the sequencing of the wild type gene of Klen Taq. Right, histogram of the nucleotide in which the autocorrelation decays to 1/e. Values over 10 are not shown.

### Error correction

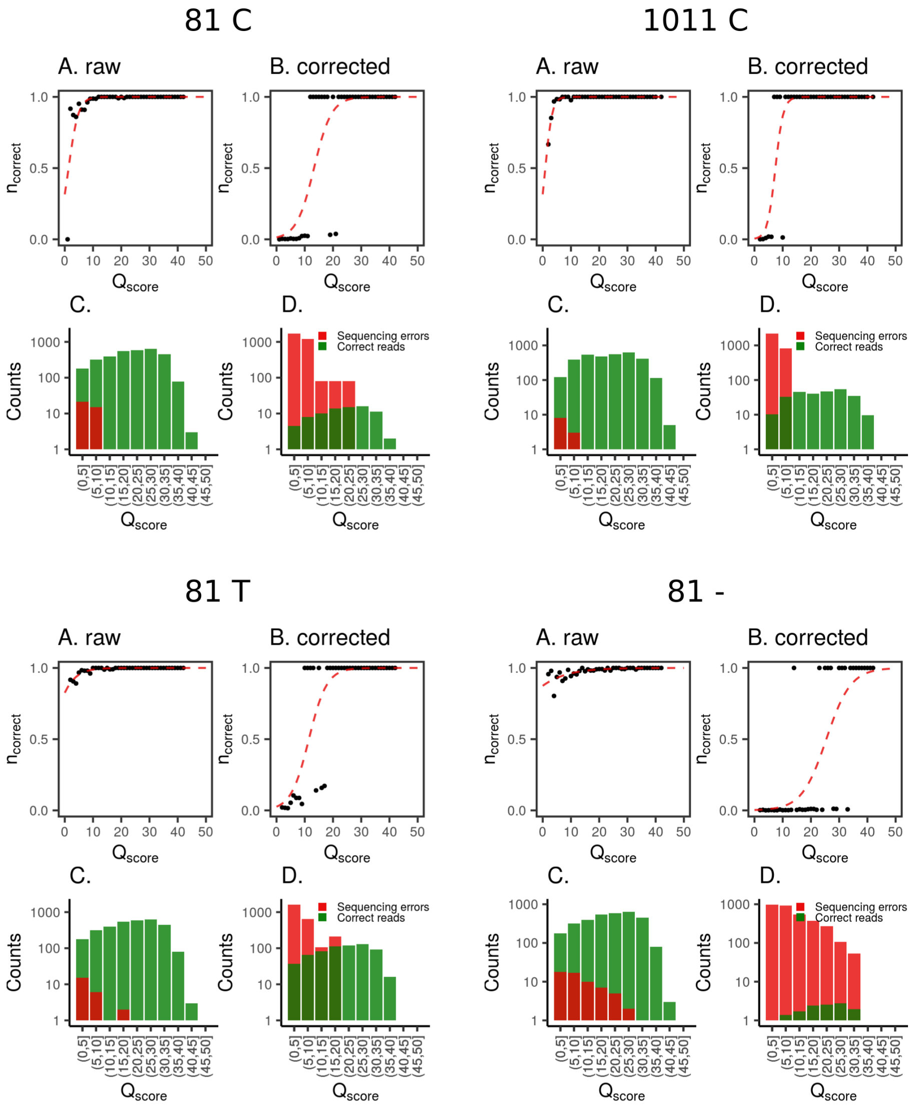


**Figure S4:** Examples of logistic fits for re-scaling of confidence values ($Q_{score}$) for different positions and nucleotides, indicated at the top. For each of them: A: black dots are the proportion of correct reads against the $Q_{score}$ reported by minION. Red dashed line is the logistic regression performed over this data. C: Distribution of correct reads (green) and sequencing errors (red). B and D: We repeat the plots A and C respectively when each read is weighted according to the prior probability of minION/guppy making an error in this position and the probability of having an actual mutation according to the ePCR kit.

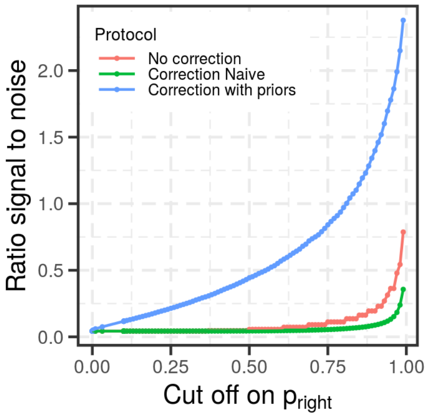


**Figure S5:** Signal to noise ratio, without weighting the counts.

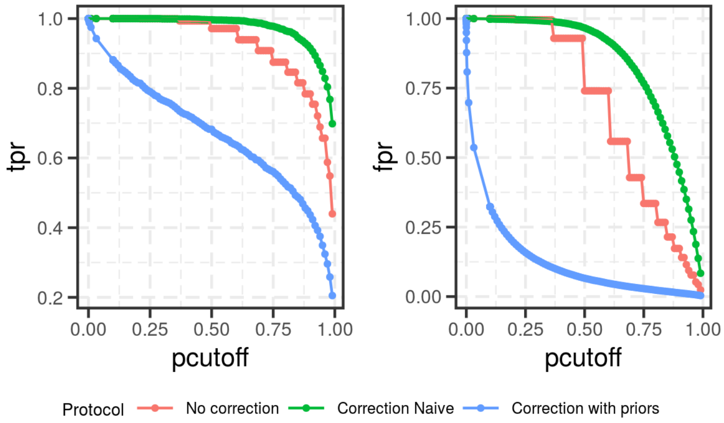


**Figure S6:** True positive rate (left) and false positive rate (right) according to the cut-off on the quality values $p_{right}$, for the three methods analyzed: using $p_{right}$ provided by guppy base-caller (red), using $p_{right}$ after naive correction (green) and $p_{right}$ after correction with priors (blue).

#### Discussion on consensus calculation

To compute the consensus sequence we used a simple computation: in an aligned set of sequences we counted at each position the occurrence of each nucleotide or deletion, weighted by $Q_{score}$ provided by guppy base-caller or the corrected $Q_{score}$ presented in the manuscript, and kept the nucleotide with higher value. Nevertheless, in the case of homopolymers, this calculation distinguishes between multiple states which are in fact equivalent. For example, if the original sequence is 'AAA' and there is a deletion, after aligning this state might appear as 'AA-', 'A-A', or '-AA'. If this alignment is not consistent in all sequences (which is the case for LAST alignments) the deletion will not be over-represented in one position and neither in the consensus (see figure S7 for a schema). In our experimental data, variant 5 represents this situation, having a deletion in a position corresponding to the homopolymer 'GG' in the wild type. It can be seen in fig. S8 (solid lines) that there is no convergence to a perfect consensus, whether we use guppy base-caller $Q_{score}$ weights or the corrected weight presented in this manuscript. Figure S9 also shows the mean number of errors per sequences is over higher in that case.
To overcome this limitation we tested two strategies (Fig. S7). In the first one, we sorted each homopolymer so that deletions are always at the end. I.e. in our example, all different states ('AA-', 'A-A', and '-AA') are replaced by the sorted state 'AA-'. Surprisingly, this had a detrimental effect in the consensus calculated with guppy base-caller $Q_{score}$ weights (Fig S8), not only for variant 5 with a deletion in a homopolymer, but for all mutants analyzed. Accumulating the deletions at the end of the homopolymers lead to a higher detection of deletions even in non mutated homopolymers (due to sequencing artifacts), whereas, without sorting, these "false" delections were averaged out among several positions. This problem is solved when deletions are re-weighted using the correction proposed, leading to a reduced number of wrong mutations in variant 5 and not disturbing the other variants significantly.
The second strategy considered each homopolymer as a unit. In our example, we do not treat each position of the homoplymer 'AAA' as a separate unit but the three as a whole. We count how many times each three-nucleotides combination appears combined and keep the most frequent one (Fig. S7). The weight are assigned as the product of the individual weights. This correction is not detrimental for the guppy base-caller $Q_{score}$ weights nor the corrected weights, while it improves the detection of the deletion in variant 5.

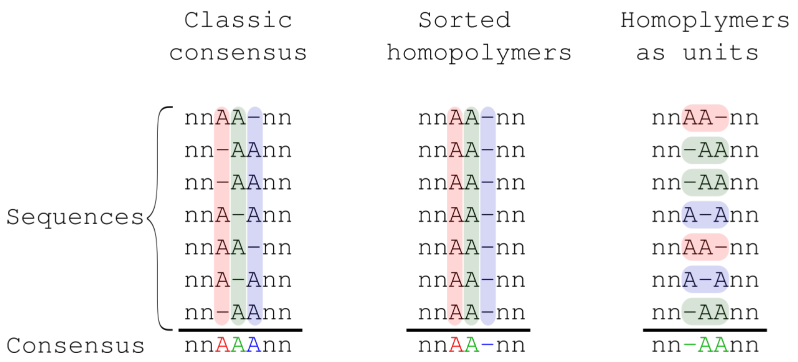


**Figure S7:** Schema of consensus calculation. Classic consensus: at each position the most frequent nucleotide is chosen. Sorted homopolymers: homopolymers are sorted to have deletions at the end. Then, at each position the most frequent nucleotide is chosen. Homopolymers as units: each homopolymer is treated as a unit, and the most frequent combination of nucleotides and deletions is chosen.

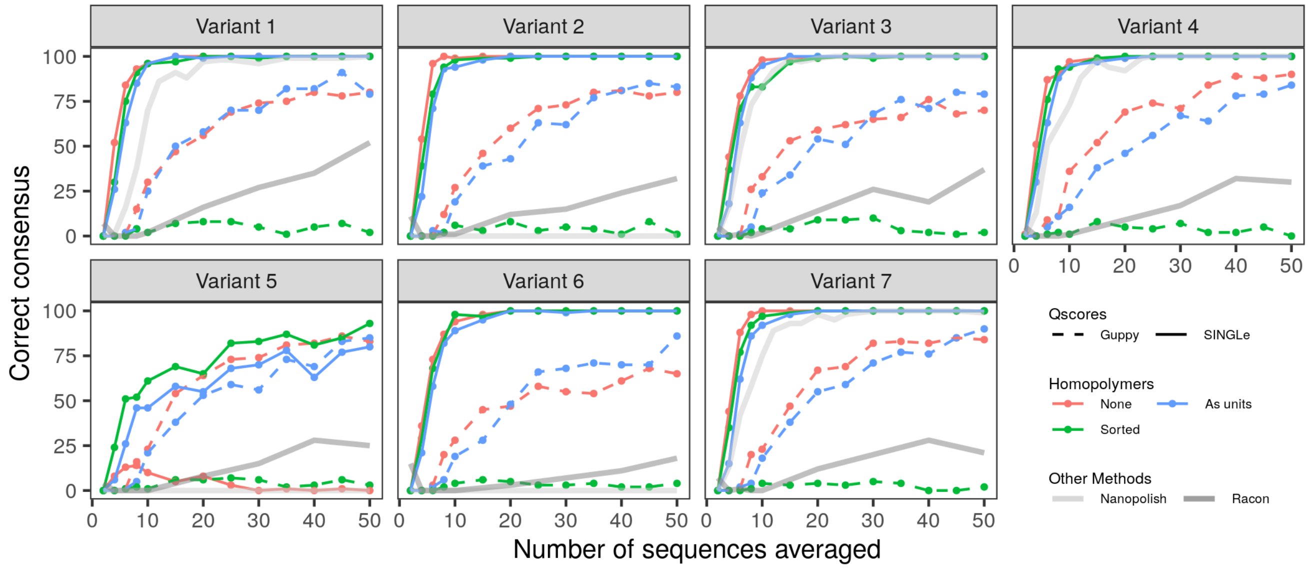


**Figure S8:** For each of the known sequenced variants (see table S2), we calculated the consensus sequence using different number of read sequences. Consensus is calculated by summing at each position the guppy base-caller $Q_{score}$ weights of the reads (red curves) or the corrected weights (black curves), and keeping the nucleotide with higher value. We draw N sequences randomly (x axis) and calculated the consensus. We repeated this calculation 100 times. In the y axis we indicate how many times the consensus matched exactly the actual sequence. The lines refer to the different consensus strategies tested, schematized in figure S7.

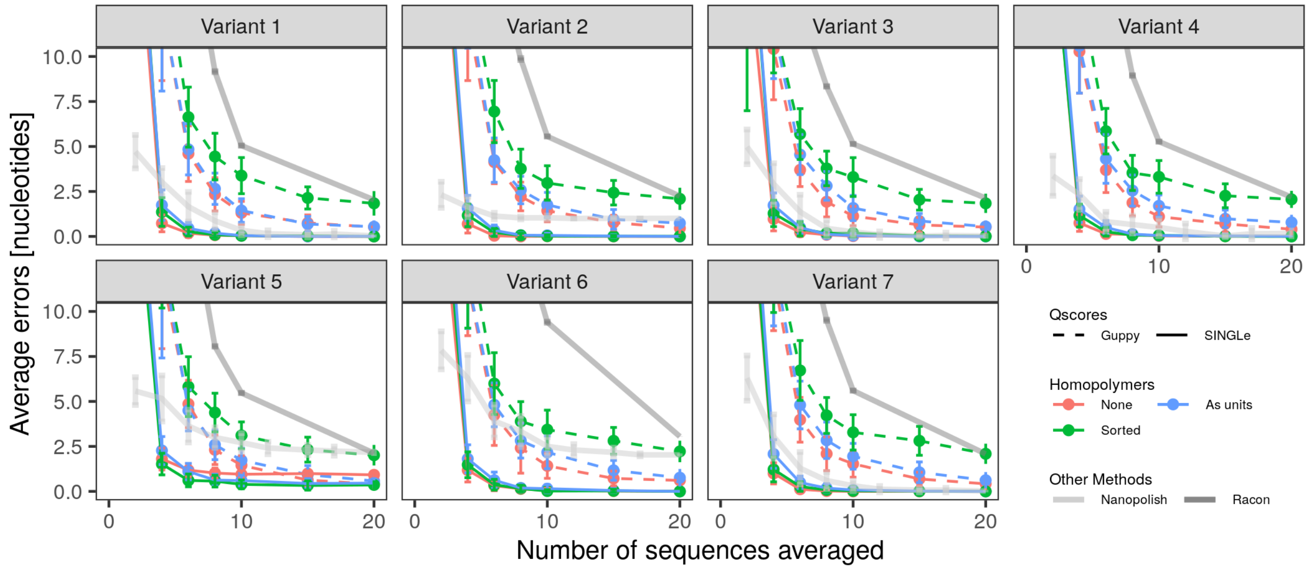


**Figure S9:** For each of the known sequenced variants (see table S2), we calculated the consensus sequence using different number of read sequences. Consensus is calculated by summing at each position guppy base-caller weights of the reads (red curves) or the corrected weights (black curves), and keeping the nucleotide with higher value. We draw N sequences randomly (x axis), calculated the consensus, and then the number of differences between the consensus and the actual sequence. We repeated this calculation 100 times for each data point, and plotted the mean number of errors and the standard deviation. The lines refer to the different consensus strategies tested, schematized in figure S7.

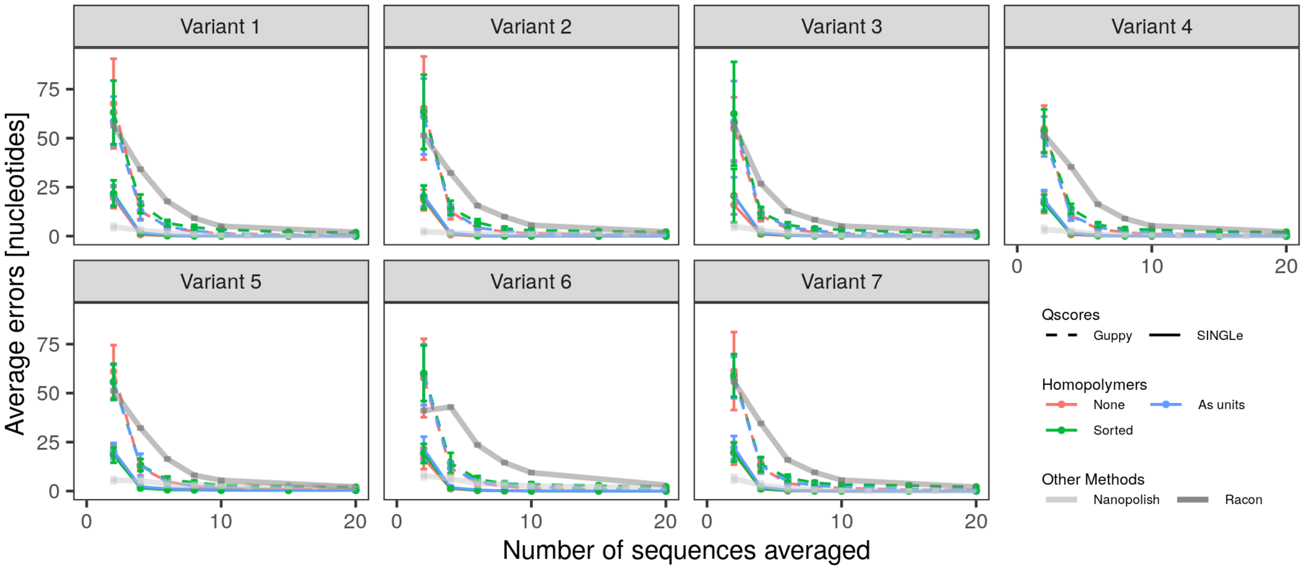


**Figure S10:** Same as figure S9, full y-range.

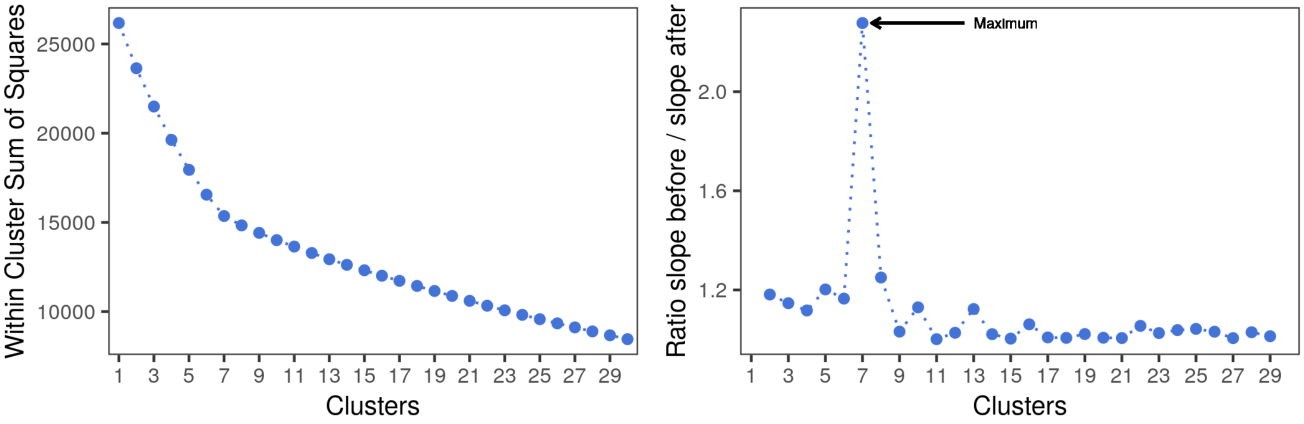


**Figure S11:** Within cluster sum of squares method to choose the number of clusters in hierarchical clustering. It is a measure of the mean squared distance between elements belonging to the same cluster. Left, within cluster sum of squares vs. number of clusters. The slope of the curve changes at 7 clusters. Right, given $s_k$ the slope between k-1 and k clusters, the change in slope is computed as $s_k s_{k+1}$. The maximum change is given at 7 clusters.
